# Splicing factor SRSF1 expands the regulatory logic of microRNA expression

**DOI:** 10.1101/2020.05.12.092270

**Authors:** Marija Dargyte, Julia Philipp, Christina D. Palka, Michael D. Stone, Jeremy R. Sanford

## Abstract

The serine and arginine-rich splicing factor SRSF1 is an evolutionarily conserved, essential pre-mRNA splicing factor. Through a global protein-RNA interaction survey we discovered SRSF1 binding sites 25-50nt upstream from hundreds of pre-miRNAs. Using primary miRNA-10b as a model we demonstrate that SRSF1 directly regulates microRNA biogenesis both *in vitro* and *in vivo*. Selective 2’ hydroxyl acylation analyzed by primer extension (SHAPE) defined a structured RNA element located upstream of the precursor miRNA-10b stem loop. Our data support a model where SRSF1 promotes initial steps of microRNA biogenesis by relieving the repressive effects of *cis*-regulatory elements within the leader sequence.

## Introduction

MicroRNAs (miRNAs) are important regulators of post-transcriptional gene expression. Nearly 60% of human protein coding genes contain conserved miRNA target sites (Friedman et al. 2009). Given the importance of miRNAs in gene regulation, it is not surprising that spatial and temporal expression patterns of miRNAs are tightly regulated. Canonical miRNA biogenesis begins with transcription of a primary miRNA (pri-miRNA) by RNA polymerase II. In the nucleus, the pri-miRNA folds into a hairpin structure which is excised by the Microprocessor complex consisting of Drosha and DGCR8, yielding a precursor miRNA (pre-miRNA). Upon transport to the cytoplasm the hairpin is cleaved, by Dicer, into a 22nt miRNA duplex. The less thermodynamically stable strand is preferentially loaded into RISC by catalytic Argonaute protein, Ago2 (Noland & Doudna 2013). Although the major catalytic steps of miRNA biogenesis and downstream RISC targeting are well understood, the regulatory checkpoints are only emerging.

RNA binding proteins are broadly implicated in miRNA biogenesis. The terminal loop region of the hairpin is a central target for many RBPs (Nussbacher & Yeo 2018; Treiber et al. 2017). For example, Lin28 binds to the terminal loop of let-7 family members recruiting TUT4 for uridylation (Heo et al. 2009). Competition between KSRP and hnRNP A1 binding to the terminal loop of pri-miR-18a influences processing by Drosha/DGCR8 (Guil & Cáceres 2007; Michlewski & Cáceres 2010). The functional importance of the terminal loop in regulation of miRNA biogenesis is underscored by strong phylogenetic conservation of this sequence element across vertebrates. In addition to the terminal loop, other sequence elements within the pri-miRNA are implicated in regulation of biogenesis (Michlewski & Caceres 2018).

The serine and arginine-rich (SR) protein family are evolutionarily conserved RNA binding proteins. Named for their Arg-/Ser-rich carboxyl terminal domain (RS domain), these proteins have diverse functions in post-transcriptional gene regulation including, pre-mRNA splicing, mRNA export, mRNA decay, nonsense mediated decay and mRNA translation (Howard & Sanford 2015). SR proteins are essential splicing factors and required for pre-mRNA splicing *in vitro* and *in vivo* (Zahler et al. 1993; Krainer et al. 1991; Li & Manley 2005). During spliceosome assembly, SR proteins, through the RS domain, promote splice site recognition via splicing factor recruitment (Zhu & Krainer 2000). Alternatively, the RS domain may function to promote RNA-RNA interactions by neutralizing electrostatic interactions between U snRNAs at the 5’ss and branch point sequence (Shen & Green 2006).

Previous work from our lab and others demonstrated that SR proteins interact with non-coding mRNA transcripts (Sanford et al. 2009; Royce-Tolland et al. 2010; Tripathi et al. 2010). By contrast to their roles in pre-mRNA splicing, the functional roles of SR proteins in small RNA expression remain poorly described. Two members of the SR protein family, SRSF1 and SRSF3, have been implicated in miRNA biogenesis. SRSF3 recognizes a sequence determinant located downstream of the basal junction in hundreds of pri-miRNAs (Kim et al. 2018; Auyeung et al. 2013). Whereas, SRSF1 promotes processing of pri-miR-7 by binding to the lower stem, although its mechanism remains unclear (Wu et al. 2010).

Here we report the discovery of a new sequence determinant of miRNA biogenesis. Using ENCODE eCLIP data, we discovered that a wide array of RBPs interact with pri-miRNAs. Remarkably, we found the region 25-50nt upstream of miRNA hairpins was a frequent ligand for RBPs, including the pre-mRNA splicing factor SRSF1. We validated these data using iCLIP, which identified hundreds of pri-miRNAs in HEK293T cells with strong cross linking signals 35-50nt upstream of the 5’ of the hairpin, which we name the 5’ leader sequence. We demonstrate that SRSF1 expression levels correlate with decreased levels of pri-miRNAs and a concomitant increase in functional miRNA activity. Using pri-miR-10b as a model, we determine that SRSF1 binding sites are necessary for SRSF1-dependent stimulation of miRNA biogenesis. Taken together our data demonstrate, for the first time, an upstream determinant required for SRSF1 directed regulation of miRNA biogenesis.

## Results and Discussion

### Global analysis of primary miRNA-protein interactions

To identify RBPs that preferentially interact with sequences outside of the hairpin, we used the ENCODE consortium enhanced crosslinking immunoprecipitation and high throughput sequencing (eCLIP-seq) data (Van Nostrand et al. 2016). We compiled more than 120 protein-RNA interactions in HepG2 and K562 cells (Supplemental Table 1). We set a range to genomic regions 100nt upstream and 200nt downstream of the 5’ end of pre-miRNAs, as annotated by Gencode. Using aggregated eCLIP peaks for all RBPs in the ENCODE database, we observed a wide array of interactions across pri-miRNAs, including a prominent region near the terminal loop region (Fig. 1B). We also noted pronounced, but broadly distributed binding sites upstream of the 5’ end of pre-miRNA. (Fig. 1B) To determine how specific RBPs interact with pri-miRNAs we plotted the binding site density for individual RBPs, with binding sites in at least 7 unique miRNAs. Unsupervised hierarchical clustering revealed that different RBPs preferentially associate with specific regions of pri-miRNAs (Fig. 1A). For example, Lin28B interacts specifically with a region encompassing the terminal loop, a finding that is well-aligned with previous studies (Choudhury & Michlewski 2012). By contrast, we noted several splicing factors, including SRSF1 and U2AF1, with preferential binding sequences upstream of the pre-miRNA (Fig. 1A and Supplemental Table 2).

**Figure 1.**
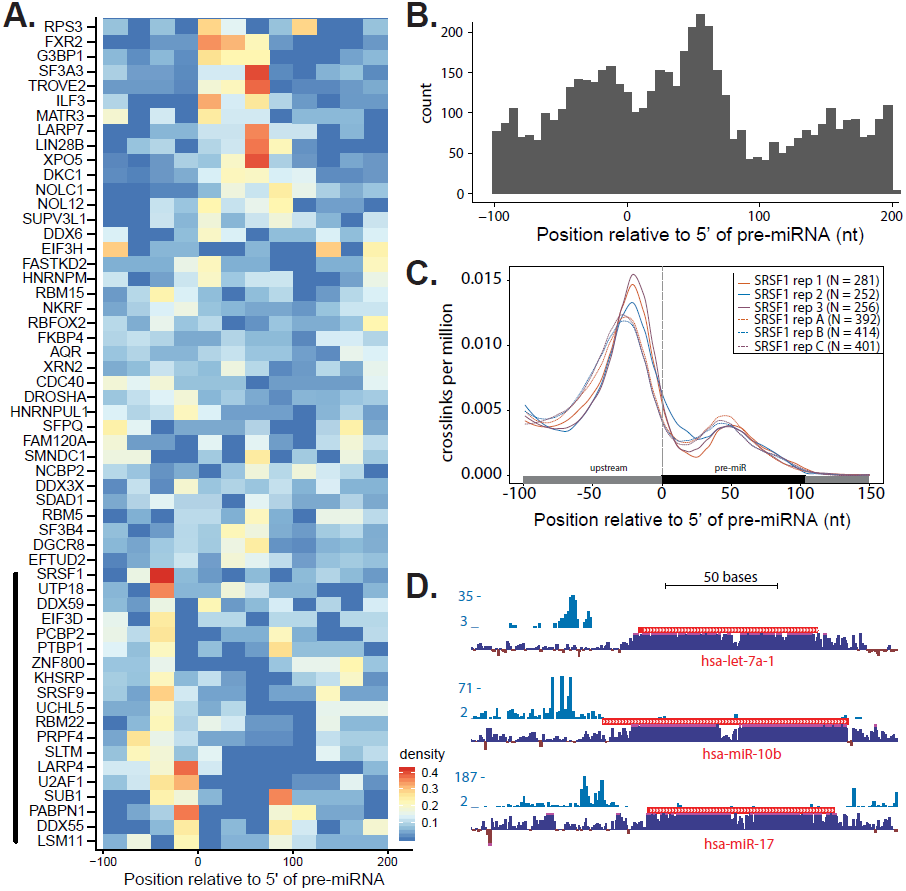
Meta analysis of eCLIP data and iCLIP data characterizes a relationship between RBP binding and pri-miRNAs. (A) Heatmap depicting specific RBP interactions along a subset of pri-miRNA transcripts using HepG2 eCLIP data. Horizontal axis denotes distance from the 5’ end of miRNAs by bins in 25nt. (B) Histogram of all RBP localizations in HepG2 cells relative to pri-miRNAs. 0 denotes 5’ end of pre-miRNAs annotated in the UCSC genome browser. (C) SRSF1 iCLIP crosslinks density relative to pri-miRNA for six replicates under two conditions. (D) UCSC genome browser screenshots of three exemplar pre-miRNAs with SRSF1 binding. Blue histogram is SRSF1 crosslinking density. Red track is pre-miRNA genes. Purple histogram is 100 vertebrate conservation.

Using published CLIP-seq and iCLIP experiments from our lab we validated the interaction of SRSF1 and the 5’ end of pre-miRNAs (Howard et al. 2018; Sanford et al. 2009). As expected, most SRSF1 binding sites identified by CLIPper in protein coding genes were associated with exonic sequences (Supplemental Fig. S1C). We also observed a purine-rich motif enriched in sequences corresponding to SRSF1 binding sites (Supplemental Fig. S1D). At a single nucleotide resolution, crosslinking density was significantly higher in exon than intron sequences, consistent with previous studies (Supplemental Fig. S1E; Sanford et al. 2009; Sanford et al. 2008; Änkö et al. 2012). We used the 5’ end of SRSF1 iCLIP reads to approximate the crosslinking position of SRSF1 on hundreds of pri-miRNAs (Supplemental Table 2). In agreement with eCLIP data, we observed a non-uniform distribution of SRSF1 crosslinking density relative to the 5’ end of pre-miRNAs, with a strong bias to positions ∼50nt upstream of the 5’ end of the pre-miRNA (Fig. 1C). SRSF1 was previously linked to regulation of miRNA processing, although the mechanism was not described (Wu et al. 2010). A curious finding from the prior study was that SRSF1 recognized a consensus binding motif located in the basal region of the pre-miR-7 stem loop. By contrast, eCLIP and iCLIP show SRSF1 interacts with sequences upstream of pre-miRNAs.

### SRSF1 stimulates miRNA activity

Our observations from both eCLIP and iCLIP data suggest that SRSF1 could be involved in miRNA regulation. To determine if SRSF1 has a global impact on miRNA expression, we sequenced small RNAs from control or SRSF1 overexpression cells. Of the 524 mature miRNAs expressed in HEK293T cells, we identified 25 upregulated and 21 are downregulated (Supplemental Fig. S2A and Supplemental Table 4). A significant majority of the differentially expressed miRNAs, ∼80% (37/46), were also detected in SRSF1 iCLIP experiments, suggesting they may be directly regulated by SRSF1. To further investigate the role of SRSF1 in miRNA expression we focused on let-7a-1, miR-10b, and -17, which stand out for their robust SRSF1 crosslinking (Fig. 1D). We measured changes in pri-miRNA levels after SRSF1 overexpression in HEK293T cells by RT-qPCR. Upon SRSF1 overexpression we observed a significant reduction in expression for each pri-miRNA as compared to control cells (Fig. 2A). Taken together, these data suggest that SRSF1 either reduces the steady state levels of pri-miRNAs via enhanced processing or RNA decay, or that SRSF1 may promote pri-miRNA processing.

**Figure 2.**
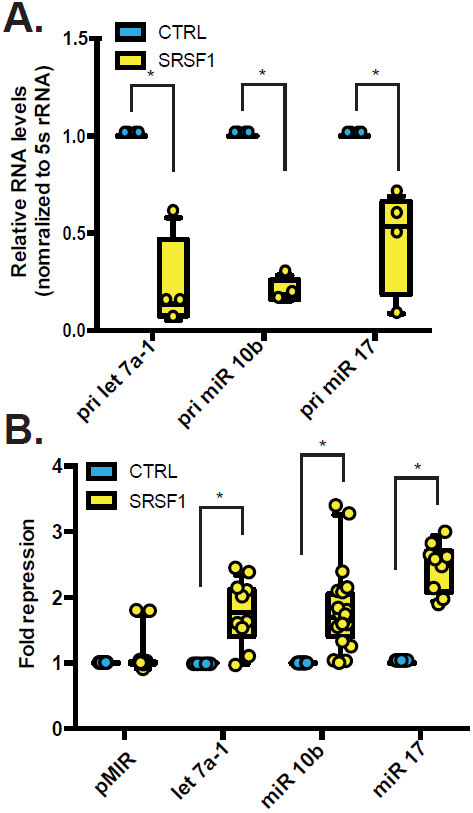
SRSF1 dependent miRNA expression and activity. (A) Relative quantification of RT-qPCR of pri-miRNAs let 7a-1, miR-10b, and miR-17. (B) Normalized luciferase reporter expression (fold repression) for let 7a-1, miR-10b, and miR-17 reporters for control (cyan) or SRSF1 overexpression (yellow). (*) P <0.05 using unpaired t-test.

To discriminate between these two hypotheses, we asked if SRSF1 overexpression influenced mature miRNA activity. We generated luciferase reporters containing target sites for specific miRNAs within their 3’UTR. Individual miRNA reporter constructs or a control reporter lacking the heterologous miRNA target site were co-transfected with T7-SRSF1 or a control plasmid into HEK293T cells. If SRSF1 stimulates either mature miRNA activity or expression we expect to see a decrease in reporter activity or an increase in repression. In all cases, we observed significant reduction in reporter activity relative to controls upon T7-SRSF1 overexpression (Fig. 2B). These data suggest that SRSF1 promotes maturation of miRNAs rather than simply reducing pri-miRNA levels. To determine if these changes in reporter activity are specific to SRSF1 we also co-transfected HEK293T cells with the same reporter constructs as well as hnRNPA1, another RBP linked to the biogenesis of specific miRNAs. As expected, over-expression of hnRNPA1 enhanced miR-17 activity (Kooshapur et al. 2018). By contrast, hnRNPA1 had no effect on let-7-a1 or miR-10b reporter activity (Supplemental Fig. S3B).

SRSF1 shuttles continuously from the nucleus to the cytoplasm and is intimately involved in mRNA processing, stability and translation (Das & Krainer 2014). To determine if SRSF1 influences a nuclear or cytoplasmic step in miRNA biogenesis we co-transfected luciferase reporters with wild type SRSF1 or a non-shuttling mutant that is retained in the nucleus (Cazalla et al. 2002). If SRSF1 promotes pre-miRNA export from the nucleus or Dicer activity in the cytoplasm, then we predict that the non-shuttling mutant would be unable to stimulate miRNA activity. By contrast, we observed that relative to wild type, the non-shuttling mutant (SRSF1-NRS) exhibits enhanced repression of the miR-10b reporter (Supplemental Fig. S3C). These data suggest that SRSF1 promotes a nuclear step in the miRNA biogenesis pathway, as previously suggested by the processing of miR-7 (Wu et al. 2010).

### SRSF1 binding sites are required for enhanced miR-10b activity in vivo

iCLIP revealed SRSF1 interactions with pri-miRNA 5’ leader sequences at single nucleotide resolution. To determine if these points of interaction are functionally relevant for miRNA processing, we generated a series of pri-miR-10b expression constructs containing point mutations at SRSF1 crosslinking sites. If SRSF1 directly promotes miRNA biogenesis, then we predict that mutation of SRSF1 interaction sites could attenuate the effect of SRSF1 on miRNA activity and expression. As expected, driving pri-miR-10b expression up in HEK293T cells strongly reduced luciferase activity relative to the negative control expression construct (Fig. 3A). Overexpression of SRSF1 further reduced miR-10b luciferase reporter activity. By contrast, pri-miR-10b expression constructs containing crosslinking site mutant 2 attenuated the effect of SRSF1 on miR-10b luciferase reporters. Similarly we observe a loss of detectable mature miR-10b with mutant 2 overexpression compared to wild type pri-miR-10b (Fig. 3B). Likewise, we do not observe significant luciferase repression changes between wild type or mutant pri-miR-10b for control experiments lacking SRSF1 overexpression (Fig. 3A). Taken together this experiment reveals at least one *cis*-acting RNA element recognized by SRSF1 functions in regulation of miR-10b expression.

**Figure 3.**
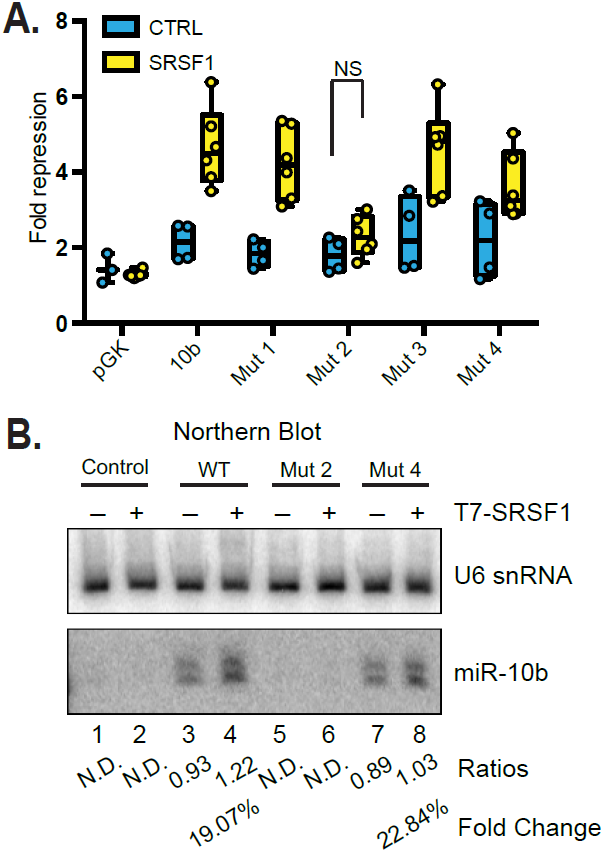
Mutations within SRSF1 binding site alter miR-10b expression and activity. (A) Normalized luciferase reporter expression (fold repression) for miR-10b reporter for control plasmid (cyan) or SRSF1 overexpression cells(yellow). (B) Northern blot of U6 snRNA (control) and mature miR-10b with overexpression of exogenous pri-miR-10b constructs along with SRSF1.

To determine if crosslinking site mutations interfere with SRSF1 pri-miR-10b interactions, we performed filter binding assays using purified recombinant SRSF1 (rSRSF1) and RNA binding-deficient mutants (Supplemental Fig. S4A). Recombinant SRSF1 binds pri-miR-10b with an apparent Kd of 31.64nM (Fig. 5A and C). As expected, rSRSF1 harboring point mutations of two solvent exposed phenylalanines in RRM1 are mutated to aspartates (FF->DD) reduce affinity for RNA binding (Supplemental Fig. S5A). Likewise, deletion of the RS domain reduces RNA binding. As expected, the point mutation in pri-miR-10b which attenuates SRSF1-dependent regulation of miR-10b activity and expression weakens the affinity of SRSF1 for pri-miR-10b *in vitro* (Supplemental Fig. S5). Although affinity for the pri-miR-10b mutants is reduced *in vitro* we cannot discount any *in vivo* interactions that are not accounted for by filter binding. These data indicate that the SRSF1 RS domain is required for binding to pri-miR-10b and that mutations within the leader do reduce SRSF1 affinity for pri-miR-10b.

### Identification of a repressive element in the 5’ leader of pri-miR-10b

The experiments described above indicate that sequences beyond the hairpin regulate pri-miRNA processing. To test this hypothesis, we created a series of deletion mutants from the 5’ or 3’ of pri-miR-10b (Fig. 4). If either the 5’ or 3’ flanking sequences are required for mature miRNA activity, we expect an increase in miR-10b luciferase reporter activity. If the mutations remove repressive elements, we expect a decrease in luciferase activity. To distinguish between these possibilities we co-transfected expression constructs for wild type pri-miR-10b or 5’ or 3’ deletion mutants, along with the miR-10b luciferase reporter. We observed significant decrease in luciferase activity for the more extreme 5’d2 and 5’d3 mutants, but not the more conservative 5’d1 mutant (Fig. 4B). These data suggest that there are sequence or structural repressive elements within the SRSF1 binding sites 5’ of the hairpin.

**Figure 4.**
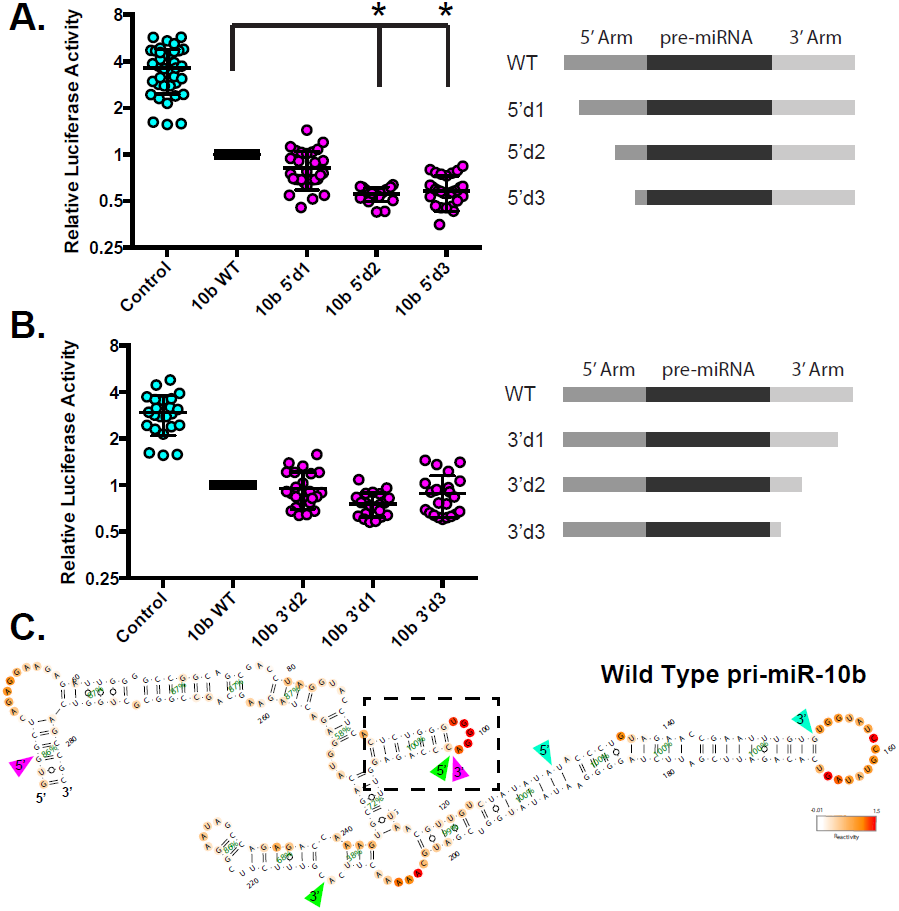
An upstream structure of pri-miR-10b influences mature miR-10b activity. (A,B) Relative luciferase activity when HEK293T cells are transfected with exogenous pri-miR-10b truncation mutations. Schematic depicts regions of truncations relative to pre-miR-10b. (*) P <0.05 using unpaired t-test. (C) Chemical probing of pri-miR-10b by 1M7. Cyan arrows denote the parameter of embedded mature miR-10b, green arrows denote the parameter of embedded pre-miR-10b, and pink arrows denote the parameter of SRSF1 crosslinking region from iCLIP. Nucleotide accessibility to 1M7 are marked by warmer colors. Note boxed, the presence of a small and stable hairpin upstream of the 5’ apical stem of miR-10b.

**Figure 5.**
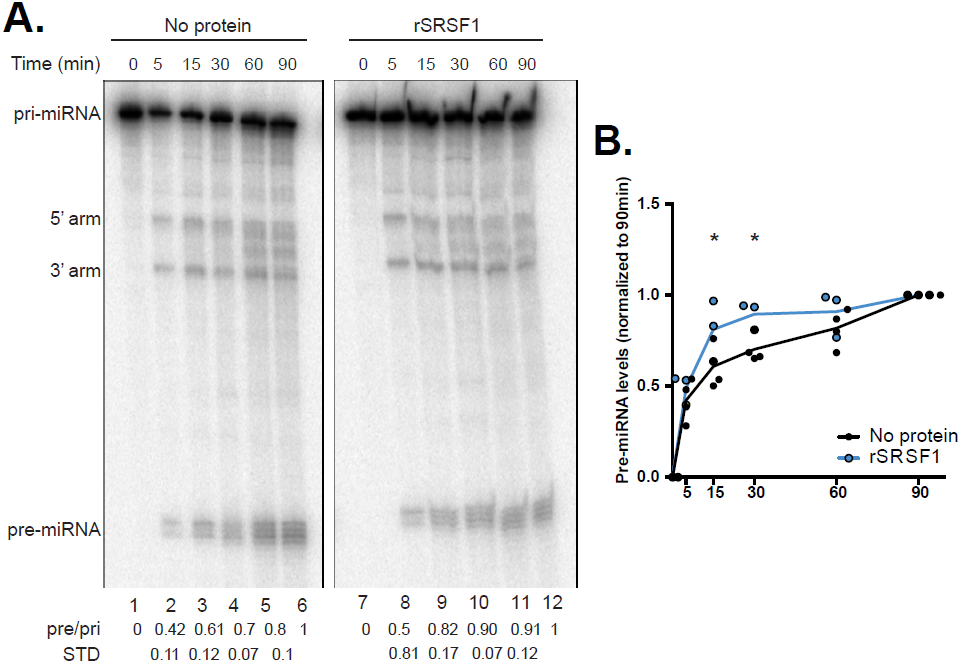
SRSF1 directly alters the rate of pri-miRNA processing. (A) In vitro processing time course of pri-miR-10b constructs by FLAG pulldown Microprocessor complex in presence or absence of rSRSF1. 5’ or 3’ arms cleaved during processing are labeled. Pri-to pre-miR-10b ratios are calculated for three replicate experiments. (B) Quantification of pre-miR-10b accumulation over 90 minutes. In presence of SRSF1 (blue) pre-miR-10b accumulated significantly faster starting at 15 minutes.

To determine if the 5’ leader of pri-miR-10b contains structured RNA elements we performed chemical probing using 1-methyl-7-nitroisatoic anhydride (1M7) SHAPE reagent. 1M7 modifies the 2’ hydroxyl of unpaired residues. Modified ribose residues are revealed as termination sites by primer extension. Using reactive positions to constrain secondary structure predictions reveals the presence of a canonical miR-10b hairpin containing the embedded mature miRNA (Fig. 4C). Surprisingly, a well defined stem loop structure emerges just upstream of the hairpin (Fig. 4C). By contrast, the point mutations that reduce the effect of SRSF1 on miR-10b activity significantly reduce reactivity within the loop region of this novel structural element, suggesting a change in secondary structure of the pri-miR-10b leader sequence (Supplemental Fig. S6).

To determine if a structured 5’ leader was a general feature of pri-miRNAs bound by SRSF1, we compared the thermodynamic stability of pri-miRNAs predicted to be bound by SRSF1 to those lacking iCLIP signal signal. Using the DINAmelt web server application, Quikfold, we were able to generate -□G values for predicted secondary structures of pri-miRNAs (Markham & Zuker 2005). We observed a slight, yet significant difference in the distribution of -□G between those primary miRNAs bound by SRSF1 and those that are not (Supplemental Fig. S7A). These data suggest that perhaps there is a structured element within the 5’ leader sequence of SRSF1 bound pri-miRNAs.

### SRSF1 directly influences miRNA biogenesis

Taken together, our results suggest that SRSF1 promotes a nuclear step of miRNA processing, and likely before initial cleavage by Drosha. Therefore we reasoned that SRSF1 may enhance Microprocessor complex activity. To test if SRSF1 directly influences the Microprocessor step of miRNA biogenesis we performed *in vitro* miRNA processing assays with immunopurified Drosha/DGCR8 in the presence or absence of rSRSF1 (Fig. 5A). In control reactions without rSRSF1 we observed a gradual increase in product formation over the course of the reaction (Fig. 5A, lanes 1-6). However, when pri-miR-10b was incubated in the presence of rSRSF1 we observed a significant increase in the rate of product formation (Fig. 5A, lanes 7-10). Quantification of replicate experiments revealed that SRSF1 enhances rates of product formation compared to control reactions (Fig. 5B). Because SRSF1 promotes pri-miR-10b processing by Drosha/DGCR8 and that Drosha contains an RS domain, it is possible that SRSF1 recruits Drosha to the pri-miRNA transcript. To test if SRSF1 directly interacts with the Microprocessor complex we probed proteins coprecipitated with Drosha by western blot. We were unable to observe any RNA-dependent or -independent interactions between exogenously expressed SRSF1 and the Microprocessor complex (Supplemental Fig. S7C). Overall, our data suggests that SRSF1 promotes pri-miRNA biogenesis by altering the conformation of the 5’ leader sequence prior to Drosha cleavage.

In this study we showed that the SR protein SRSF1 promotes the first steps in miRNA processing. Global analysis of protein-RNA interactions by iCLIP and eCLIP revealed that SRSF1, as well as other splicing factors, engage binding sites upstream of pre-miRNAs (Figure 1). Reporter assays demonstrated that SRSF1 enhances miRNA function *in vivo* and that *cis*-acting SRSF1 binding sites within pri-miR-10b are required. Our data suggests that this 5’ leader sequence is inhibitory, and needs to be relieved for efficient processing. Alleviating a repressive domain for miRNA biogenesis has been previously described and well supported by our data as well (Du et al. 2015). This observation is strongly supported by *in vitro* processing assays, which show that rSRSF1 accelerated the cleavage rate of pri-miR-10b. Coimmunoprecipitation experiments failed to detect an interaction between SRSF1 and Drosha, arguing against a recruitment model. Instead, we suggest that SRSF1 may influence the conformation of the pri-miRNA. Using SHAPE we noted the presence of a strong stem loop structure within the 5’ leader region of primary miR-10b. Deletion analysis suggests the 5’ leader region interferes with miR-10b expression. Taken together our data suggest that SRSF1 binding to pri-miR-10b alters the conformation of an inhibitory stem loop structure.

Despite decades of research, the mechanisms through which SR proteins regulate post-transcriptional gene expression remain unclear. Competing models include RS domain recruitment of splicing factors and RNA-RNA interaction chaperones (Graveley & Maniatis 1998; Shen & Green 2006). Previously, ATP-independent RNA annealing activity was copurified with SRSF1 (Krainer et al. 1990), suggesting that SRSF1 disrupted intramolecular RNA structure formation to promote intermolecular annealing at temperatures well below the T_m_. One prediction is that SRSF1 relieves inhibitory secondary structures in the 5’ leader sequence. We believe such a mechanism is consistent with our observations using pri-miR-10b as a model. This structural change could serve as a checkpoint in hairpin selection by the Microprocessor. A similar licensing step was described for processing of the pri-miR-17∼92 cluster (Du et al. 2015).

The results presented here, demonstrate that SRSF1 promotes miRNA processing without directly recruiting the Microprocessor. Given the recent discovery that SRSF3 influences miRNA processing through interactions with the basal junction (Kim et al. 2018). We hypothesize that SRSF1 and SRSF3 may function collaboratively, by 5’ and 3’ interactions respectively, to define the hairpin for miRNA processing. This process likely involves remodeling inhibitory secondary structure adjacent to the stem loop and consistent with an RNA chaperone function for SRSF1 in miRNA biogenesis.

## Materials and Methods

### Analysis of eCLIP and iCLIP datasets

eCLIP data was downloaded from the ENCODE consortium through their dashboard. Peak definitions from HEPG2 cells were aligned relative to the 5’ end of miRNA precursors. Data were visualized following unsupervised hierarchical clustering. Only RBPs with at least 7 annotated binding sites near miRNAs were considered in this analysis (Supplemental Table 1 and 2). iCLIP data for SRSF1 was downloaded from (GSE #GSE83923). Reproducible crosslinking sites were defined as previously described (Howard et al. 2018). Crosslinking density was calculated for all SRSF1 crosslinking data relative to the 5’end of miRNA precursors.

### Cell culture and transfections

Hek293T cells were grown in 6 well plates with DMEM supplemented with 10% FBS. At 70% confluence cells were transfected with plasmids using polyethylenimine (PEI) and 0.35M NaCl. Each transfection was performed a minimum of three times with two technical replicates per experiment.

### RNA purification and RT-qPCR

Total RNA for RT-qPCR was isolated using Direct-zol RNA MiniPrep Kit (Zymo Research) for all other experiments RNA was isolated using standard Tri-reagent (Sigma) protocol. cDNA was reverse transcribed from 1ug of total RNA using High-Capacity cDNA reverse transcriptase kit (Applied Biosystems). qPCR was performed using Titanium Taq (Clontech) and SYBR Green on a Roche Lightcycler 480 (Roche Diagnostics) according to MIQE guidelines (Bustin et al. 2009).

### Luciferase reporter assays

Seed sites for let-7a-1, miR-15b, miRNA17, miR 19a, miR 93, and miR 128a were inserted into the 3’UTR of pMIR luciferase reporter (Life Scientific). miR-10b reporters described previously (Ma et al. 2007) were obtained from AddGene. Reporters were co-transfected with Renilla luciferase (Promega) reporter as a transfection efficiency control. Luciferase activity was assayed 24 hours post transfection using Dual-Glo Luciferase Assay System (Promega). For a 24-well plate, each well was transfected with 100ng of TK-rLUC (Promega), 800ng or 1ug of T7-SRSF1 or control plasmid (Cáceres et al. 1997), 400ng of pMIR Luciferase reporter (Life Scientific). Experiments in which exogenous pri-miR-10b was used, cells were transfected with 200ng of pGK (control) or pGK 10b (Ma et al. 2007; Cáceres et al. 1997).

### In vitro transcription

20ug of linearized plasmid with BamHI (New England Biolabs) of which 2ug was transcribed with MEGAscript T3 polymerase (ThermoFisher). Transcripts were labeled with alpha-^32^P UTP for in vitro processing and filter binding. Following transcription, RNA was phenol/chloroform extracted and ethanol precipitated. RNA was resolved on a 6% denaturing polyacrylamide gel and extracted with a clean razor. Gel containing RNA was incubated overnight at 42°C in elution buffer (0.3M NaOAc pH 5.5, 2% SDS). RNA was ethanol precipitated and stored at -20°C until use.

### In vitro miRNA processing

FLAG-immunoprecipitate *in vitro* processing assays were performed based on (Lee et al. 2003). Briefly, HEK293T cells were transfected with equal concentrations of FLAG-Drosha and HA-DGCR8 which were FLAG immunoprecipitated according to protocol. Processing reactions were incubated as a time course up to 90 minutes at 37°C. RNA was phenol/chloroform extracted and resolved on a 10% denaturing polyacrylamide gel. After drying, gel was exposed on a phosphor screen, visualized with a Typhoon image scanner (GE Healthcare), and was analyzed using ImageJ. Data was standardized between gels by normalizing the ratio of pre-to pri-miRNA bands from the 90 minute time point.

### Northern blot

10-25ug of total RNA was resolved on a 12.5% denaturing polyacrylamide gel and transferred to a positively charged nylon membrane (GE Healthcare) using 1x TBE. After UV crosslinking, membrane was prehybridized with ULTRAhyb hybridization buffer (Invitrogen) for 30 minutes at 42°C. The membrane was hybridized to a ^32^P end labeled oligo probe overnight at 42°C. Membrane was washed with 2X SSC, 0.05% SDS, and 0.1X SSC, 0.1% SDS respectively for 30 minutes at 42°C. Blot was visualized using Typhoon image scanner (GE Healthcare) and analyzed using ImageJ.

## Acknowledgments

We thank DS Rao and JM Draper for critical comments on the manuscript. We thank Dr. Demián Cazalla for Drosha/DGCR8 constructs. We thank A. Zahler and M. Ares for thoughtful discussion. This research was supported by NIH grants GM130361 (JRS). and GM095850 (MDS).

## Author Contributions

JRS conceived the study. JRS, MD, JP, CDP, MDS. designed the experiments. MD, CDP, JP. performed the experiments. MD, CDP, JP. analyzed the data. JRS and MD. wrote the manuscript.

## Supplemental Methods

### Small RNA sequencing

For small RNA sequencing experiments total RNA was provided to RealSeq BioSciences (Santa Cruz, CA) and converted to small RNA-seq libraries using the RealSeq-AC kit. Sequencing statistics for each library are available in Supplemental Table 3.

### Recombinant SRSF1 protein purification

Purification of recombinant SRSF1 and mutant proteins was performed as previously described (Cazalla et al. 2005). Briefly, 30ug of SRSF1 expression plasmids were transfected into 15 cm^2^ plates of HEK293T cells using Lipofectamine 2000 (ThermoFisher). 48 hours post transfection cells were washed in ice cold PBS and lysed in 50 mM NaP buffer, pH 8, 0.5 M NaCl, 5 mM β-glycerophosphate, 5 mM KF, 0.1% Tween 20, and 1× protease inhibitor cocktail. T7-Tag Agarose Beads (Novagen) was used to affinity purify T7-tagged SRSF1 from cleared lysates as previously described.

Proteins were dialyzed overnight in BC100 (20 mM Tris, pH 8, 100 mM KCl, 0.2 mM EDTA pH 8, and 20% glycerol). Protein concentrations were quantified by A280 reading on Nanodrop and BCA assays (ThermoFisher).

### Filter binding assays

0.25 nM of body labeled RNA was heated for 3 minute at 90°C in 50mM HEPES and cooled to room temperature for 15 minutes. RNA was incubated with increasing concentrations of protein and 10mM MgCl_2_ for 30 minutes at 37°C. Reaction mixture was then loaded into a 96-well vacuum minifold dot blot (Whatman) using a house vacuum and washed with 1mL of 50mM HEPES (pH 8.0) and 10mM MgCl_2_. Protein bound RNA was collected on nitrocellulose (Amersham) and unbound RNA was collected on positively charged nylon membrane (Ambion). Blots were exposed overnight on a phosphor screen, and lane analysis of dot intensities was performed using imageJ. Background was obtained in the absence of protein and subtracted from the data set.

### Western Blot Analysis

Proteins were separated on 10% SDS-PAGE, transferred onto a nitrocellulose membrane (Bio-rad) and blotted against T7 (Millipore), SRSF1 (SCBT), HSP90 (SCBT) or EWS (SCBT). Signal was detected using SuperSignal West Pico PLUS Chemiluminescent Substrate (ThermoFisher).

## Supplemental figure legend

**Supplemental Figure S1.** SRSF1 iCLIP results. (A) Autoradiograph of protein-RNA complexes. Black line denotes where within the smear in lane 4, protein-RNA complexes were excised. (B) Pie chart denoting where SRSF1 iCLIP peaks map back to the genome. (C) Consensus motif for SRSF1 derived from above iCLIP. (D) Graph depicting SRSF1 crosslinking sites relative to intron-exon junctions. Note SRSF1 crosslinks higher over exons. Blue line is SRSF1 crosslinks from SRSF1 overexpression background, while red line is SRSF1 crosslinks from a control cell line.

**Supplemental Figure S2.** Small RNA (smRNA) sequencing reveals changes in miRNA levels post SRSF1 overexpression. (A) Western blot confirming T7-SRSF1 overexpression in triplicate in samples used for smRNA sequencing. (B) Heat map showing miRNAs that are up (red) and down (blue) regulated after SRSF1 overexpression. (C) Venn diagram depicting degree of overlap between miRNAs that are differentially expressed from smRNA-seq and iCLIP.

**Supplemental Figure S3.** (A) Luciferase reporter activity for let-7a-1, miR -10b, -15b, - 17, -19a, -93, -128a for control (cyan) and SRSF1 overexpression (yellow). (B) Luciferase reporter activity for let-7a-1, miR -9, -10b, -15b, -17 for control (cyan) and hnRNP A1 overexpression (brown). (C) Luciferase reporter activity for miR-10b for control (cyan), SRSF1 (yellow), and SRSF1-NRS mutant (orange) overexpression.

**Supplemental Figure S4.** Purification of T7-SRSF1 and mutants from HEK293T cells. Commasie stained gels from SRSF1 (A), SRSF1-FFDD (B), and (C) SRSF1 delRS. From left to right L is lysate, F is flow through, W is wash, 1-10 are 10 serial elutions. Two of the highest intensity elutions were collected for dialysis.

**Supplemental Figure S5.** Quantified filter binding assay measuring the fraction of 0.25nM RNA bound by protein. (A) Wild type pri-miR-10b binding to SRSF1 (brown) or SRSF1 mutants, FFDD (orange) and delRS (green). (B) Wild type SRSF1 binding to wild type pri-miR-10b (brown), or pri-miR-10b mutants; mutant 2 (orange), or mutant 4 (brown). (C) Table lists relative Kds for wild type or mutant pri-miR-10b binding to wild type SRSF1 or mutant SRSF1. Mutants have very low to no binding to RNAs resulting in large Kds.

**Supplemental Figure S6.** Secondary structures of pri-miR-10b by 1m7 chemical probing. Structures for wild type pri-miR-10b (A), mutant 4 (B), and mutant 2 (C). Arrows distinguish important features of the pri-miRNAs. SRSF1 cross linking parameter (purple), annotated pre-miRNA (green), and embedded mature miR-10b (cyan). Red arrows denote sites of substitution mutations for the individual mutations. Note boxed, the presence of a small and stable hairpin upstream of the 5’ apical stem of miR-10b. The color bar denotes nucleotide reactivity, red being most solvent accessible. Structures were aligned by orientation of their terminal loop.

**Supplemental Figure S7.** Predicted global RNA structures and protein-RNA interactions. (A) Predicted Gibbs free energies calculated for pri-miRNAs either bound or not bound by SRSF1, as defined by iCLIP. (*) P <0.05 using unpaired t-test. (B) Western blots for T7, Drosha, and DGCR8 from T7 immunoprecipitation. Control or T7-SRSF1 overexpressing cells were immunoprecipitated followed by RNase digestion (lanes 5-8). (C) Protein-protein interactions for Drosha, DGCR8, and SRSF1 attained from BioGRID.

## Notes

### Competing Interest Statement

The authors have declared no competing interest.

