## Supplementary material for "Splicing factor SRSF1 expands the regulatory logic of microRNA expression": Sup Fig 1

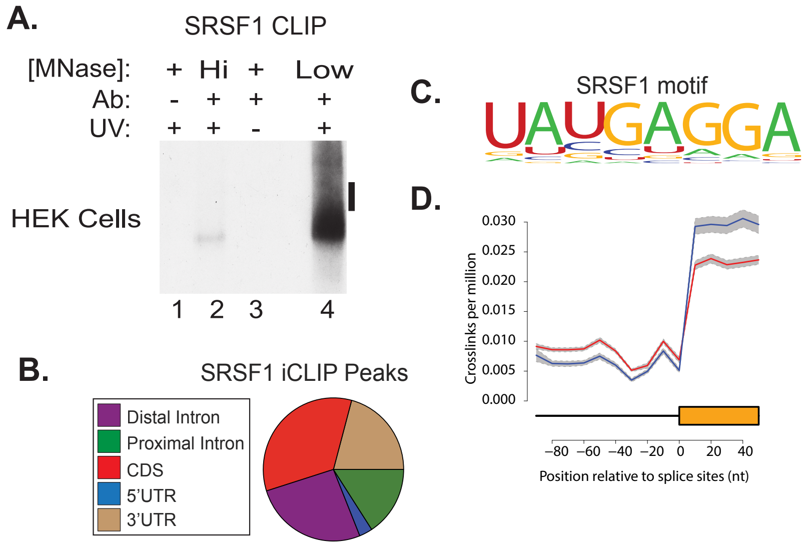


**Supplemental Figure S1.** SRSF1 iCLIP results. (A) Autoradiograph of protein-RNA complexes. Black line denotes where within the smear in lane 4, protein-RNA complexes were excised. (B) Pie chart denoting where SRSF1 iCLIP peaks map back to the genome. (C) Consensus motif for SRSF1 derived from above iCLIP. (D) Graph depicting SRSF1 crosslinking sites relative to intron-exon junctions. Note SRSF1 crosslinks higher over exons. Blue line is SRSF1 crosslinks from SRSF1 overexpression background, while red line is SRSF1 crosslinks from a control cell line.
