## Supplementary material for "Splicing factor SRSF1 expands the regulatory logic of microRNA expression": Sup Fig 2

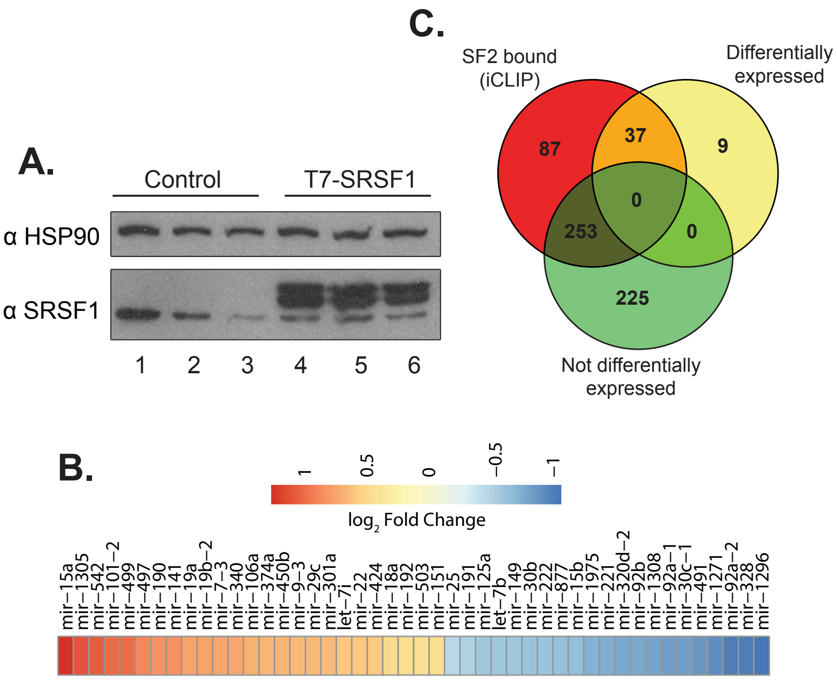


**Supplemental Figure S2.** Small RNA (smRNA) sequencing reveals changes in miRNA levels post SRSF1 overexpression. (A) Western blot confirming T7-SRSF1 overexpression in triplicate in samples used for smRNA sequencing. (B) Heat map showing miRNAs that are up (red) and down (blue) regulated after SRSF1 overexpression. (C) Venn diagram depicting degree of overlap between miRNAs that are differentially expressed from smRNA-seq and iCLIP.
