## Supplementary material for "Splicing factor SRSF1 expands the regulatory logic of microRNA expression": Sup Fig 3

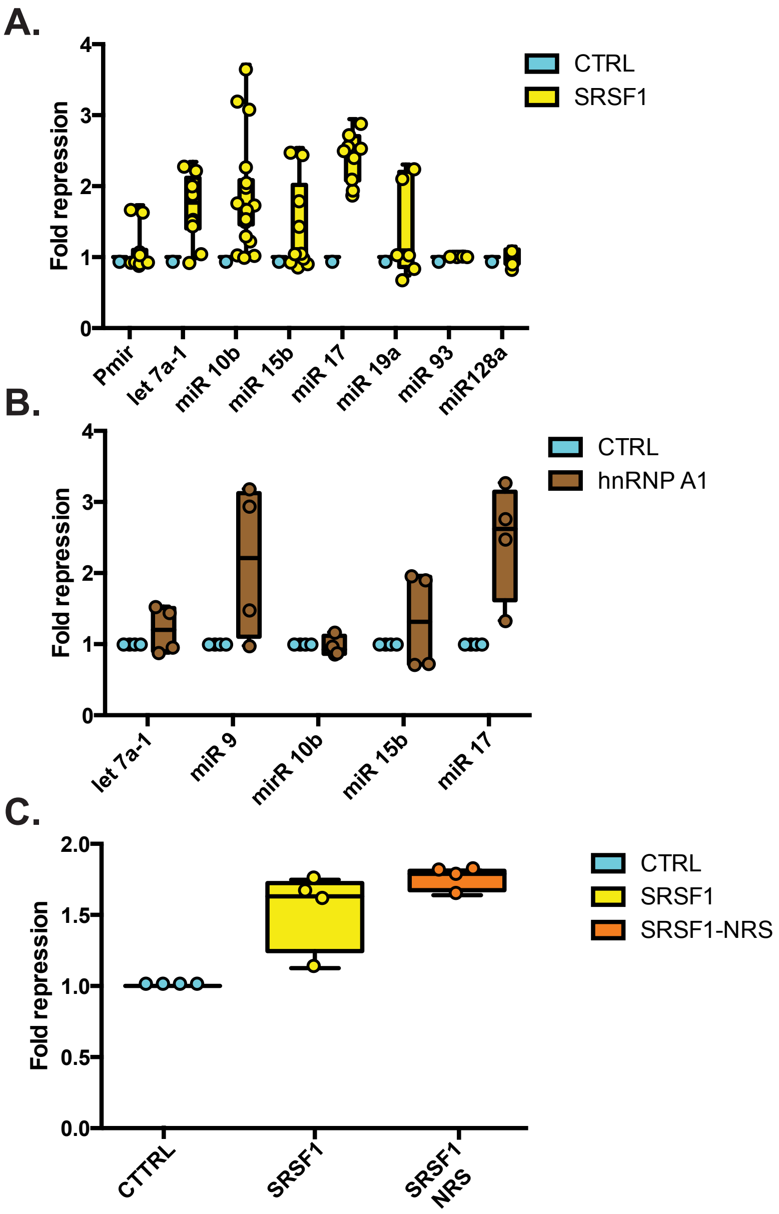


**Supplemental Figure S3.** (A) Luciferase reporter activity for let-7a-1, miR -10b, -15b, -17, -19a, -93, -128a for control (cyan) and SRSF1 overexpression (yellow). (B) Luciferase reporter activity for let-7a-1, miR -9, -10b, -15b, -17 for control (cyan) and hnRNP A1 overexpression (brown). (C ) Luciferase reporter activity for miR-10b for control (cyan), SRSF1 (yellow), and SRSF1-NRS mutant (orange) overexpression.
