## Supplementary material for "Splicing factor SRSF1 expands the regulatory logic of microRNA expression": Sup Fig 4

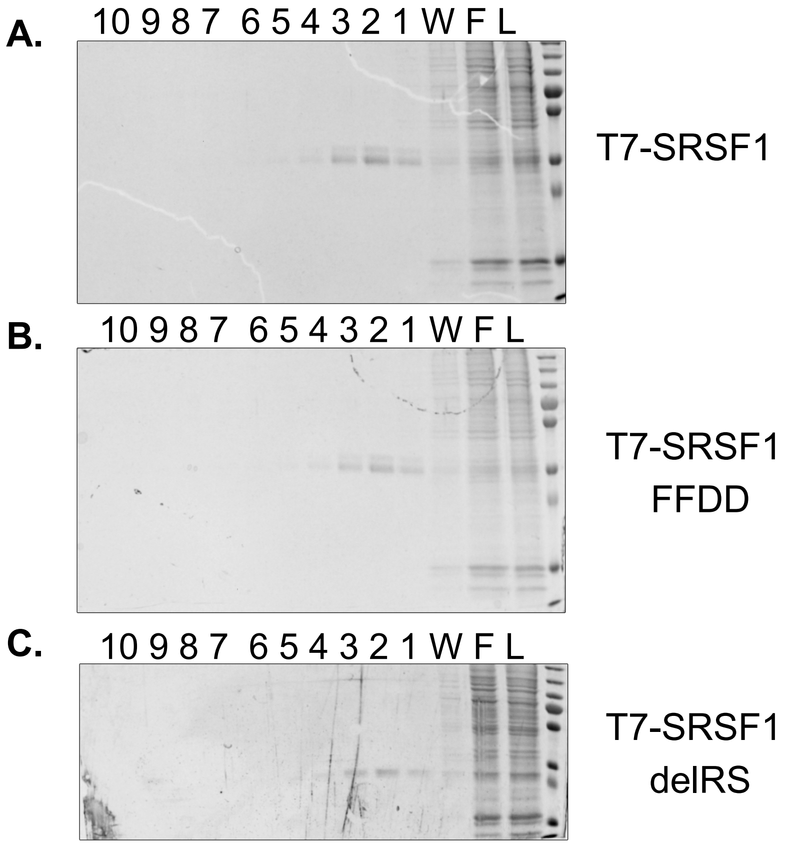


**Supplemental Figure S4.** Purification of T7-SRSF1 and mutants from HEK293T cells. Commasie stained gels from SRSF1 (A), SRSF1-FFDD (B), and (C) SRSF1 delRS. From left to right L is lysate, F is flow through, W is wash, 1-10 are 10 serial elutions. Two of the highest intensity elutions were collected for dialysis.
