## Supplementary material for "Splicing factor SRSF1 expands the regulatory logic of microRNA expression": Sup Fig 5

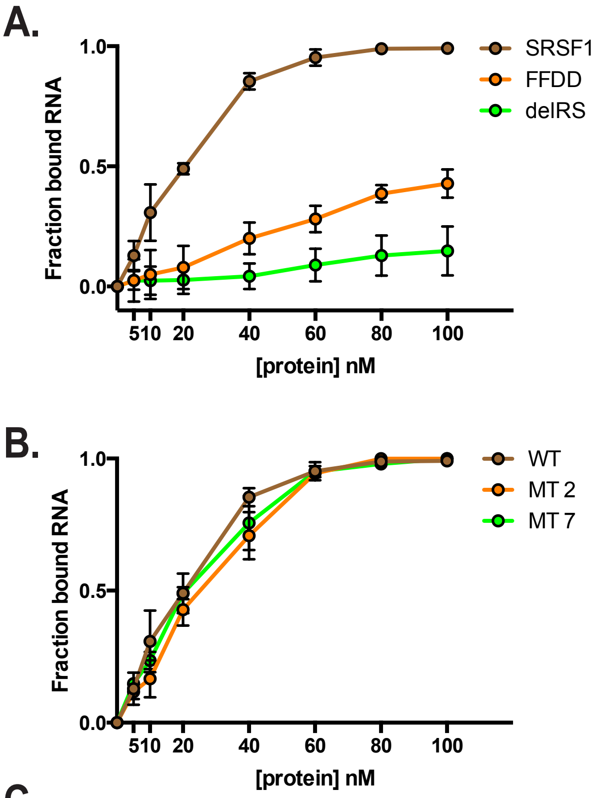


C.

| Dissociation constants determined by filter binding assay (nM) | | | |
|---|---|---|---|
| | **SRSF1** | **SRSF1 FFDD** | **SRSF1 del RS** |
| **pri-miR-10b WT** | 31.64 +/- 2.5 | 555.64 +/- 532 | N.D. |
| **pri-miR-10b MT 2** | 57.54 +/- 5.2 | 182.8 +/- 64 | N.D. |
| **pri-miR-10b MT 4** | 41.28 +/- 3.0 | 174.3 +/- 83 | N.D. |

**Supplemental Figure S5.** Quantified filter binding assay measuring the fraction of 0.25nM RNA bound by protein. (A) Wild type pri-miR-10b binding to SRSF1 (brown) or SRSF1 mutants, FFDD (orange) and delRS (green). (B) Wild type SRSF1 binding to wild type pri-miR-10b (brown), or pri-miR-10b mutants; mutant 2 (orange), or mutant 4 (brown). (C) Table lists relative Kds for wild type or mutant pri-miR-10b binding to wild type SRSF1 or mutant SRSF1. Mutants have very low to no binding to RNAs resulting in large Kds.
