## Supplementary material for "Splicing factor SRSF1 expands the regulatory logic of microRNA expression": Sup Fig 6

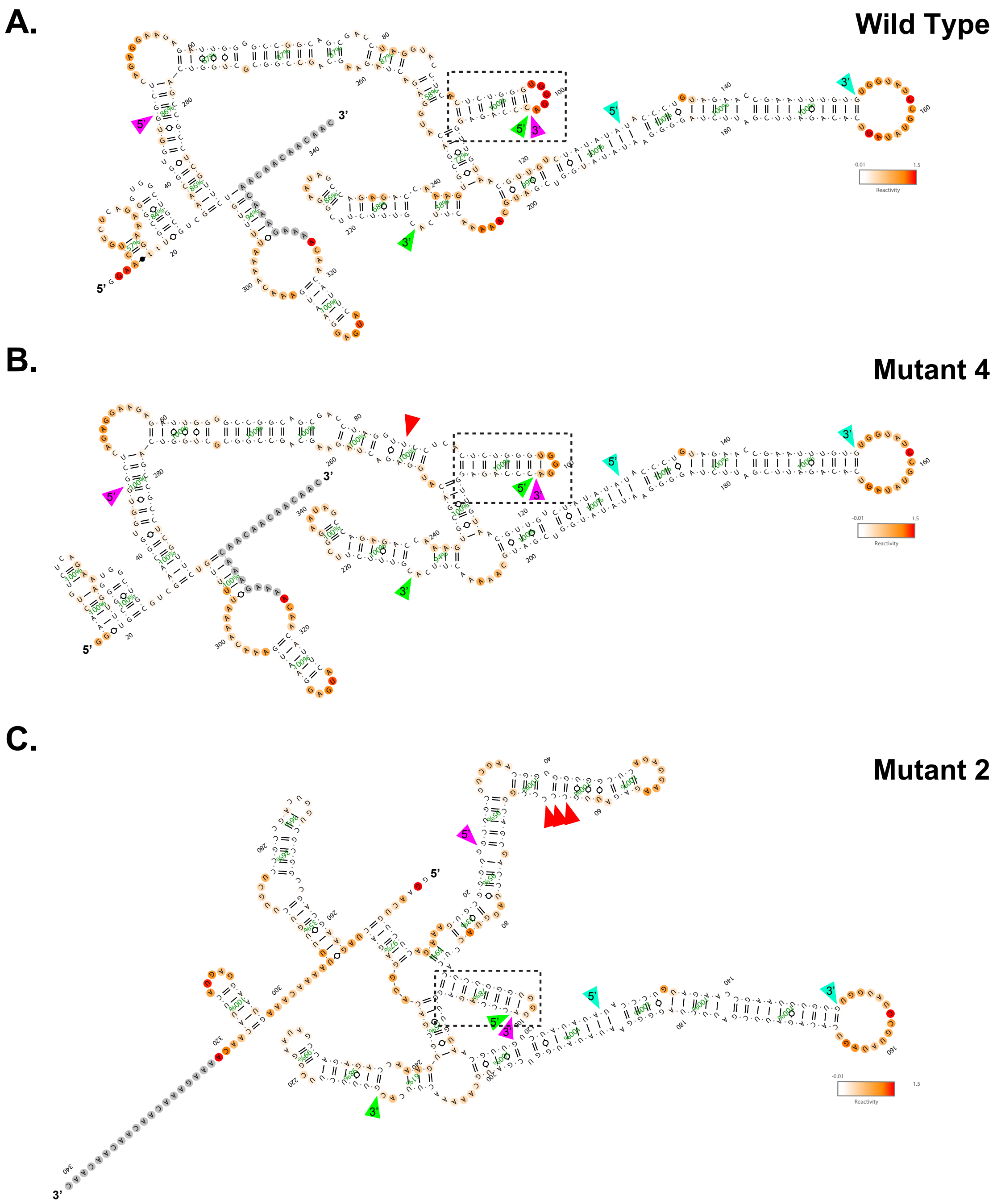


**Supplemental Figure S6.** Secondary structures of pri-miR-10b by 1m7 chemical probing. Structures for wild type pri-miR-10b (A), mutant 4 (B), and mutant 2 (C). Arrows distinguish important features of the pri-miRNAs. SRSF1 cross linking parameter (purple), annotated pre-miRNA transcript (green), and embedded mature miR-10b (cyan). Red arrows denote sites of substitution mutations for the individual mutations. Note boxed, the presence of a small and stable hairpin upstream of the 5’ apical stem of miR-10b. The color bar denotes nucleotide reactivity, red being most solvent accessible. Structures were aligned by orientation of their terminal loop.
