## Supplementary material for "Splicing factor SRSF1 expands the regulatory logic of microRNA expression": Sup Fig 7

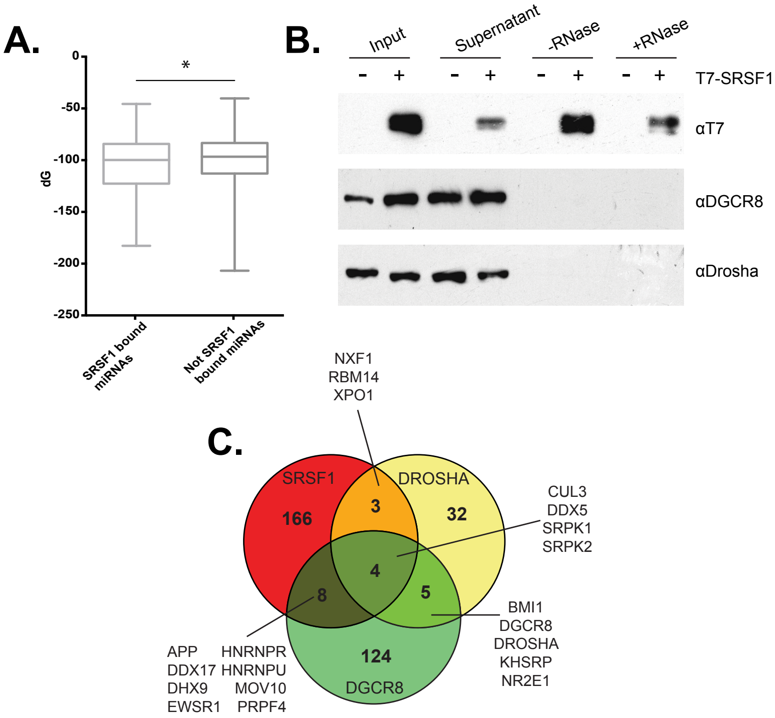


**Supplemental Figure S7.** Predicted global RNA structures and protein-RNA interactions. (A) Predicted Gibbs free energies calculated for pri-miRNAs either bound or not bound by SRSF1, as defined by iCLIP. (*) P <0.05 using unpaired t-test. (B) Western blots for T7, Drosha, and DGCR8 from T7 immunoprecipitation. Control or T7-SRSF1 overexpressing cells were immunoprecipitated followed by RNase digestion (lanes 5-8). (C) Protein-protein interactions for Drosha, DGCR8, and SRSF1 attained from BioGRID.
