## Supplementary material for "Splicing factor SRSF1 expands the regulatory logic of microRNA expression": Sup Methods

*RNA structure probing and modeling*

1.2 pmols of RNA in 50 mM Na-HEPES (pH 8) was denatured at 95℃ for 3 minutes, after cooling to room temperature, 10 mM of MgCl2 was added to the solution. RNA was modified in presence of 5 mM of 1-methyl-7-nitroisatoic anhydride (1M7, provided by Dr. Manuel Ares) at room temperature for 15 minutes, followed by quenching with 2.55 M Na-MES (pH 6) for 10 minutes. RNA was washed with 70% ethanol and purified with AMPure Beads (Beckman Coulter). After elution in water, RNA was reverse transcribed by standard SuperScript III protocol (ThermoFisher) and hydrolyzed by 200 mM NaOH.

cDNA is purified as previously described on AMPure beads and eluted with 11ul of formamide-ROX350 mix (ThermoFisher) for 15 minutes at room temperature. Primer extension products were analyzed by Elim Biopharmaceuticals by capillary electrophoresis. Data from capillary electrophoresis runs was analyzed with HiTRACE MATLAB package as previously described (Yoon et al. 2011; Palka et al. n.d.).
